# Targeting of plasmodesmal proteins requires unconventional signals

**DOI:** 10.1101/2022.11.11.516208

**Authors:** Gabriel Robles Luna, Jiefu Li, Xu Wang, Li Liao, Jung-Youn Lee

## Abstract

Cellular signaling relies on precise spatial localization and dynamic interactions of proteins within the subcellular compartment or niches, including cell-cell contact sites and connections. In plants, both endogenous and pathogenic proteins gained the ability to target plasmodesmata, the membrane-lined cytoplasmic connections, to regulate or exploit cellular signaling across cell wall boundaries. Those include the receptor-like membrane protein PDLP5, a potent regulator of plasmodesmal permeability that generates feed-forward or -back signals vital for plant immunity and root development. However, little is known about what molecular features determine the plasmodesmal association of PDLP5 or other proteins. Notably, although these proteins each have the ability to target plasmodesmata, no protein motifs or sequences have been identified indicative of targeting signals. To address this knowledge gap, we combined machine learning and targeted mutagenesis approaches. Here we report that PDLP5 and its closely related proteins carry novel targeting signals comprising short stretches of amino acid residues. As for PDLP5, it contains two non-redundant, tandemly arranged signals, either of which is sufficient for both localization and biological function regulating viral movement. Strikingly, plasmodesmal targeting signals exhibit little conservation in sequence but are located similarly proximal to the membrane. These novel unconventional features appear to be a common theme in plasmodesmal targeting.

## INTRODUCTION

Plasmodesmata form cylindrical membrane nanopores vital to allow for various nutrients and signal carrying molecules to directly move between neighboring plant cells. They are highly dynamic intercellular pores capable of undergoing rapid changes in permeability in response to signals communicating physiological and developmental state as well as mechanical, biotic and pathogenic stressors (Sager and Lee, 2014; Lee, 2015; Stahl and Faulkner, 2016; Cheval and Faulkner, 2018). At the structural level, plasmodesmal pores formed in higher plants consist of outer and inner membrane leaflets that create cytoplasmic sleeves through which soluble molecules are thought to move. The outer and inner plasmodesmal membranes are morphologically continuous to the plasma membrane (PM) and the endoplasmic reticulum (ER) membrane, respectively (Tilsner et al., 2016). However, compositionally, not all ER or PM proteins associate with plasmodesmata, raising the question of whether plasmodesmal localization may require a specific mechanism of targeting or lateral translocation.

To date, various membrane proteins have been identified to target plasmodesmata. These include single- and multi-pass integral transmembrane proteins that play key roles regulating plasmodesmal permeability, such as callose synthases, cell surface receptors, and receptor-like proteins represented by plasmodesmata-located proteins (PDLPs) (Vaten et al., 2011; Faulkner et al., 2013; Stahl et al., 2013; Cui and Lee, 2016; O’Lexy et al., 2018; Rosas-Diaz et al., 2018). Others include proteins involved in regulating cytoskeletal components or involved in processes such as membrane tethering (Vaddepalli et al., 2014; Diao et al., 2018; Brault et al., 2019). Nonetheless, how these integral membrane proteins specifically target plasmodesma (hereafter, referred to as plasmodesmal proteins) remains poorly understood.

Of known *Arabidopsis* plasmodesmal proteins, about 15 are single-pass, type-I TM proteins having an N-terminal signal peptide and a C-terminal anchoring TM domain (TMD). These include an eight-membered family of plasmodesmata located proteins (PDLPs), a few formin-like actin-binding proteins, and receptor-like kinases (RLKs) of various protein families, such as SUB1, ACR4, BAM1, and FLS2 (Faulkner et al., 2013; Stahl et al., 2013; Vaddepalli et al., 2014; Diao et al., 2018; Rosas-Diaz et al., 2018; Wang et al., 2020). PDLPs are distinct in that all members primarily target plasmodesmata while the latter proteins tend to accumulate both at the PM and plasmodesmata. Although these all associate with plasmodesmata, they do not share any bioinformatically identifiable common motifs suggestive of subcellular localization signals. However, a few were reported to require their TMDs for plasmodesmal localization (Stahl et al., 2013; Vaddepalli et al., 2014). As for the PDLP family, we recently reported that their TMDs do not ascribe targeting specificity, and foreign TMDs can substitute innate TMDs in PDLP1, PDLP5, or other PDLP members without affecting their targeting (Wang et al., 2020). Taken together, it is unclear if there are different ways that TMDs contribute to plasmodesmal localization or that the TMDs simply serve a role in membrane anchoring without being a sorting determinant.

PDLP5 is a potent plasmodesmal regulator involved in immune, physiological, and developmental signaling (Lee et al., 2011; Wang et al., 2013; Lim et al., 2016; Toyota et al., 2018; Aung et al., 2020; Sager et al., 2020; Fichman et al., 2021). It regulates plasmodesmal permeability by activating specific callose synthases to deposit callose around plasmodesmata (Cui and Lee, 2016). It localizes to plasmodesmata exclusively, which was demonstrated at the ultrastructural level by correlative light and electron microscopy and immunogold labeling (Lee et al., 2011). However, how this exclusive plasmodesmal localization is achieved remains unknown. The extracellular domain (ExD), constituting the bulk of PDLP5, contains two repeats of domain of unknown function (DUF) 26, each characterized primarily by 6 conserved cysteine residues that engage in intramolecular disulfide bond formation. Beyond this feature and an evolutionarily conserved motif within the TMD (Wang et al., 2020), PDLPs are highly variable in sequence.

In this report, we present the data showing that plasmodesmal targeting requires unconventional signals. We began our investigation using PDLP5 as a model protein for molecular dissection using a combination of molecular and cellular approaches and machine learning. We then applied the knowledge gained from dissecting PDLP5 to other plasmodesmal proteins to tease out molecular determinants required for targeting. Our experimental data indicate that an extracellular region of PDLPs, previously unrecognized as having a function common to these proteins due to its high variation in sequence among them, in fact carries bona fide <u>p</u>lasmo<u>d</u>esmal targeting signals.

## RESULTS

### A novel extracellular region carries a signal that targets PDLP5 to plasmodesmata

Having learned from our prior research that both ExD and TMD of PDLP5 do not determine localization (Wang et al., 2020), we turned our attention to its cytosolic tail for further investigation. We have previously found that a truncation of the cytosolic tail such that it leaves the TMD as the free C-terminus results in a loss of expression in either in a transient expression system using *Nicotiana benthamiana* or a stable system using transgenic *Arabidopsis* (Wang et al., 2020). To assess the importance of this sequence, we generated construct with PDLP5 lacking its cytosolic tail as a C-terminal fusion to the enhanced green fluorescent protein (EGFP) and transiently expressed the resulting fusion protein in *N. benthamiana* leaf epidermal cells.

Confocal imaging of those cells showed a normal plasmodesmal localization pattern similar to a wild-type PDLP5-EGFP, as indicated by punctate fluorescent signals along the cell boundaries, which colocalize with aniline blue-stained callose (ΔCT in Fig. 1 & Fig. S1). We conclude that the cytosolic tail sequence is dispensable for plasmodesmal localization.

**Fig. 1.**
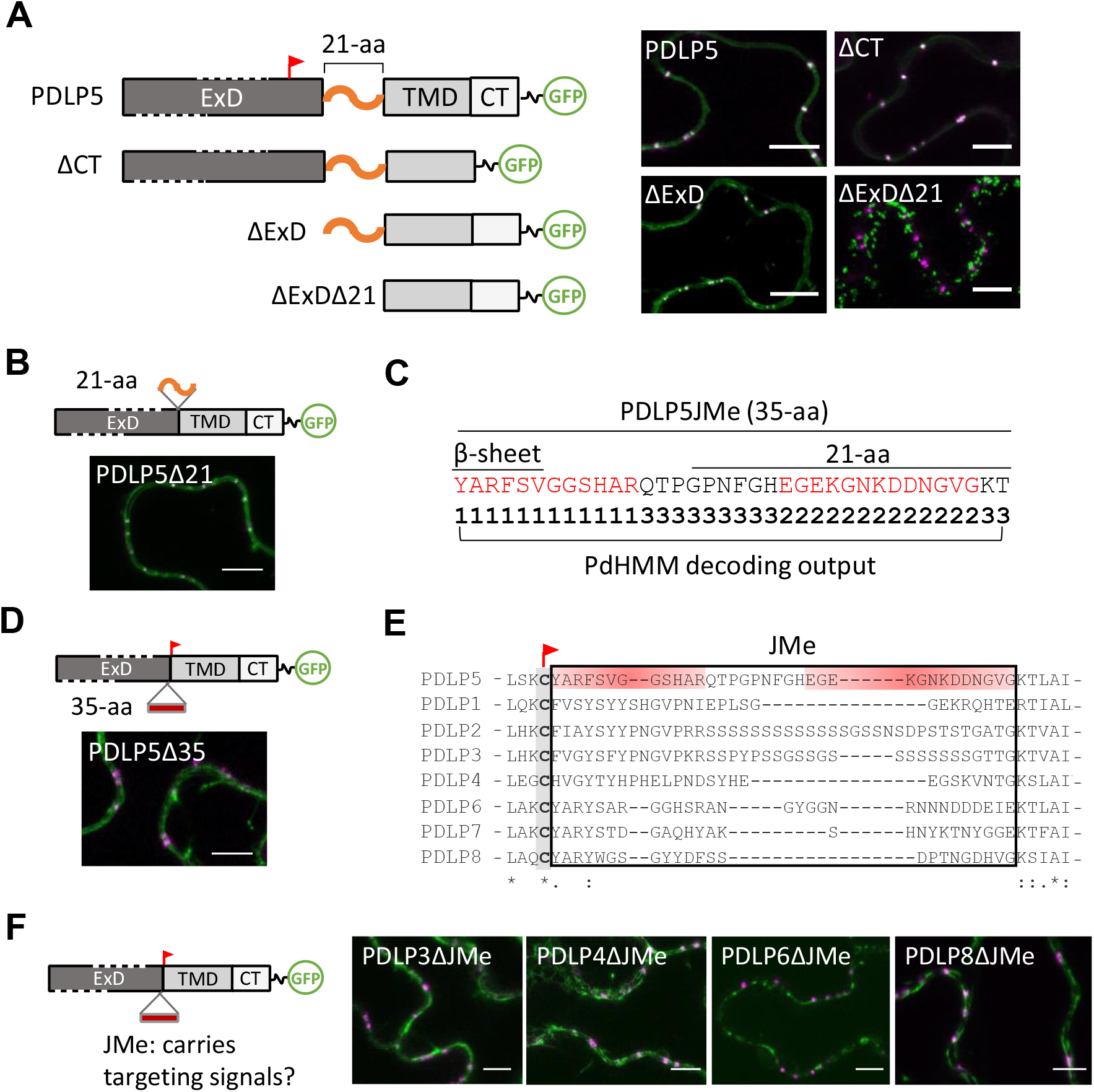
Plasmodesmal targeting requires an extracellular region. **A**. Diagrams depicting full-length and deletion mutant constructs of PDLP5 and representative confocal images showing their normal or lack of plasmodesmal localization. Dark gray boxes represent the ExD, extracellular domain consisting of tandem DUF26 modules (red flag, the last conserved cysteine in the second DUF26 module); light gray boxes, the TMD (transmembrane domain); white boxes, the CT (cytosolic tail). Orange curved lines, 21-aa presumed to be a junctional sequence. **B & D**. Representative confocal images showing subcellular localization of PDLP5 mutants lacking 21- or 35-aa. **C**. The sequence of the 35-aa region with decoding information. 1s and 2s, residues that are predicted to encode signals (colored in red); 3s, residues that are predicted to encode no signals. **E**. Sequence alignment of PDLPs showing the extracellular junctional region (JMe). Red flag, the last conserved cysteine. **F**. Subcellular localization of PDLP mutants lacking JMe sequences. **A-F**, All GFP fusion constructs were each transiently expressed in *N. benthamiana* leaf epidermal cells. Plasmodesmal localization is confirmed by co-localization of GFP fluorescence (false-colored in green) with aniline blue-stained callose (false-colored in magenta) using live-cell imaging by confocal microscopy, which is visualized as punctate whitish signals at the cellular boundaries in merged images (See figures S2, S3 and S5 for split channels). Scale bars, 10 µm.

Having exhausted all recognizable primary domains for examination, we decided to scrutinize the shortest mutated version of PLDP5 in hand, ΔExD, that maintains plasmodesmal localization (Wang et al., 2020) (Fig. 1A & Fig. S2). This sequence primarily consists of the TMD and the cytosolic tail but also includes 21-aa (amino acid) residues on the extracellular portion of the molecule. The latter sequence is located between the ExD and TMD where sequences are highly variable in both length and composition among PDLPs (Fig. S3). In addition, a predicted structure that became available from AlphaFold (Tunyasuvunakool et al., 2021) suggested that this region is likely unstructured (Fig. S3). Given that both TMD and the cytosolic tail do not determine localization, we explored if that junctional sequence is a crucial element for targeting. For this, we created a shorter version of ΔExD by deleting the relevant 21-aa. To our surprise, removing this sequence impaired plasmodesmal localization and retained the mutated version of PLDP5 at Golgi-like vesicles (Δ21 in Fig. 1A & Fig. S2). This result indicates the high likelihood that this extracellular 21-aa sequence encompasses the signal required to localize PDLP5 to plasmodesmata.

### Computational modeling predicts two targeting signals in PDLP5

Subsequently, we performed an experiment to validate the role of the extracellular sequence in the context of full-length PDLP5 by deleting the relevant 21-aa residues, but the resulting protein still retained plasmodesmal localization (PDLP5Δ21 in Fig 1B & Fig. S2). Although baffled by this result, we hypothesized the possibility of a second targeting signal located elsewhere in PDLP5. To explore this idea, we undertook a machine learning approach to identify candidate sequences. For this computational decoding task we needed to specify a segment of sequence likely to contain both targeting signals and recognize that the unknown second signal may reside near the 21-aa region. Taking these points and cumulative experimental data into account, we rationalized assigning the segment to include the 21-aa residues and residues located at their N-terminal side immediately distal to the last conserved cysteine of the second DUF26 module. We refer to this region as JMe for juxta <u>m</u>embrane sequence positioned at the <u>e</u>xtracellular side of the protein. There are 35-aa residues in total in PDLP5JMe, which includes in addition to the 21 residues, 6-aa residues that constitute the last β-sheet of the DUF26 fold (Vaattovaara et al., 2019a) followed by 9 residues comprising an unstructured region (Fig. 1C & Fig.S3).

Next, to decode the PDLP5JMe sequence, we used a hidden Markov model (HMM) because HMM can capture subtle sequence patterns by incorporating correlations of nearby residues. Since there are no clear sequence patterns in the junctional regions of PDLPs (see Fig. S3) we selected an algorithm that does not require a multiple sequence alignment for the process and built a model we named PdHMM (see Methods) Using this model, we could generate a decoding output, which predicted two stretches of sequence at the N- and C-terminal sides of the segment as signals, separated by non-signal residues (Li et al., 2020a) (Fig. 1C & Table S1). The two signal zones are similar in size, consisting of approximately 12-13 residues. Notably, the C-terminally positioned zone fell within the previously identified 21-aa region, while the N-terminally positioned zone immediately followed the last conserved cysteine among PDLPs.

With the machine-predicted decoding data in hand, we next assessed its overall validity by examining if the 35-aa sequence comprising all of JMe is required for targeting. Indeed, the mutant lacking the JMe segment was completely impaired in targeting plasmodesmata and was retained in the ER (PDLP5Δ35 in Fig. 1D & Fig. S2). This result indicates that the JMe region is vital for plasmodesmal targeting of PDLP5. It also suggests that the model may be correct in predicting the location of the two distinct, yet potentially redundant, targeting signals.

### PDLPs require the extracellular region for plasmodesmal localization

A multiple sequence comparison shows that PDLPs are highly variable in the JMe region (Fig. 1E). This type of variable region would usually be considered unlikely to harbor a domain with functional significance across the protein family. However, having learned the crucial function of JMe in PDLP5, we were interested if other PDLPs also carry the targeting signals in their corresponding JMe segments. To address this, we created mutated versions with JMe segments removed for several PDLPs, and remarkably, they were also all impaired in their subcellular localization (Fig. 1F & Fig. S4). This result suggests that the JMe region likely contains targeting signals across PDLPs despite its variability in aa sequence.

### ne targeting signal is sufficient for plasmodesmal localization

With the decoding prediction of PDLP5JMe in hand and having corroborated the significance of the JMe region for targeting, we next pursued targeted mutagenesis experiments of the two computationally predicted signals. Here, we named the two predicted signals N and C, accounting for their relative position in the sequence segment (Fig. 2A & Fig. S5). The signal mutants created using a combination of alanine substitution and truncation were then expressed transiently in *N. benthamiana* and subcellular localization followed by confocal microscopy.

**Fig. 2.**
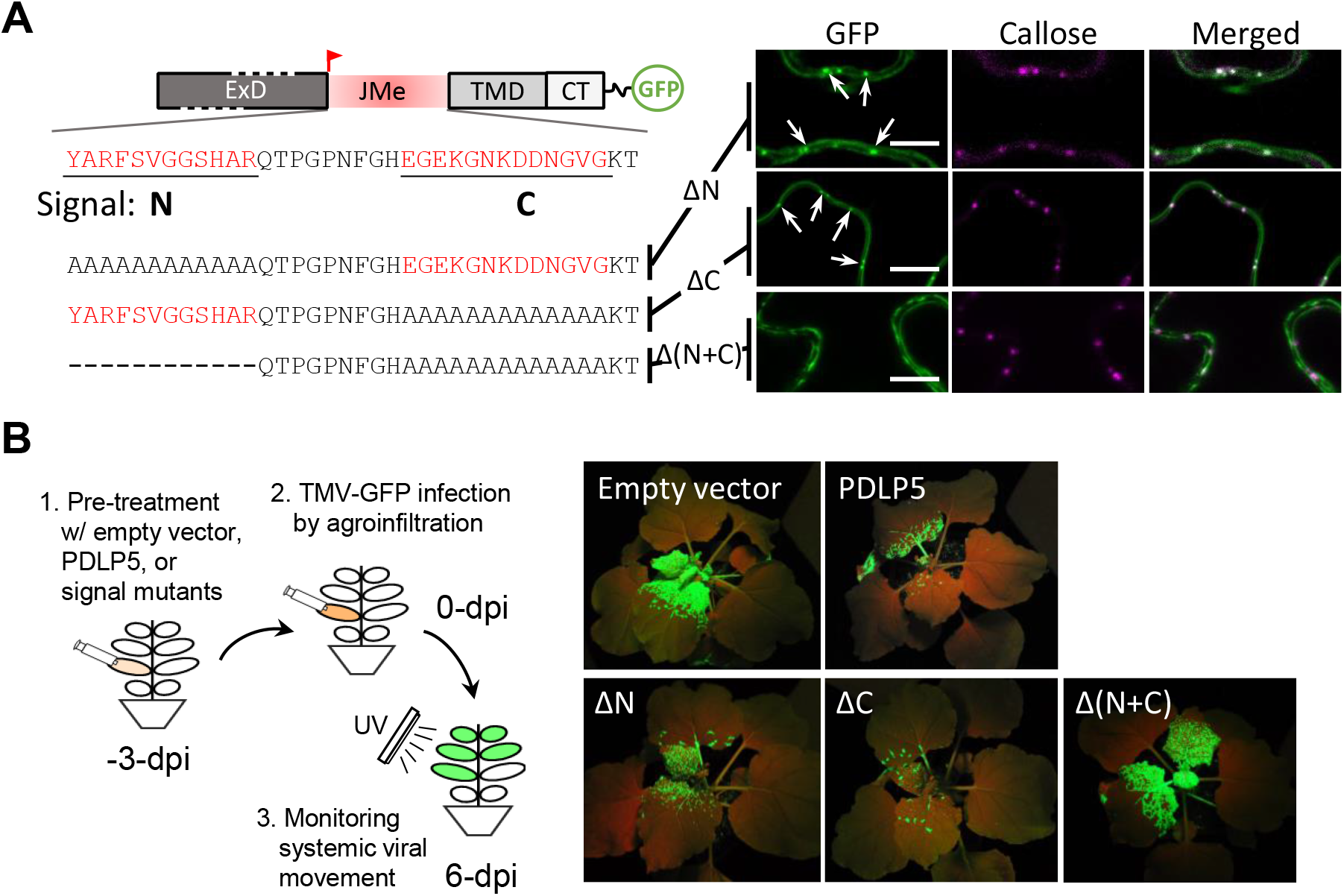
One targeting signal is enough for the normal localization and function of PDLP5. **A**. Left, a diagram showing the location and sequences of two machine-predicted signals N and C, and mutations made to eliminate them, ΔN, ΔC, and Δ(N+C). Right, representative confocal images. Arrows, plasmodesmata. Scale bars, 10 µm. **B**. Functional analyses of ΔN, ΔC, and Δ(N+C) compared with wild-type PDLP5 by viral movement assays. Left, a cartoon illustrating the experimental setup; right, representative plant photos showing the extent to which TMV-GFP moved systemically. Experiments were repeated three times using at least 5 plants per treatment.

This revealed that while either of the signals, N or C, was sufficient for plasmodesmal targeting, perturbing both simultaneously completely impaired correct targeting. These results were corroborated in transgenic *Arabidopsis* plants (Fig. S6).

### One targeting signal is sufficient for PDLP5 function

Having found that one signal is sufficient for targeting PDLP5 to plasmodesmata, we next asked if having just one signal would be sufficient for PDLP5 to exert its function. To answer this question, we examined if the versions of PDLP5 with modified signal sequences are similar or differ from native PDLP5 in their suppressive effects on viral movement in *N. benthamiana*, as described elsewhere (Lee et al., 2011; Wang et al., 2020). For this, an engineered version of *Tobacco mosaic virus* construct encoding free GFP from the viral genome (TMV-GFP) was introduced into *Agrobacterium tumefaciens* and inoculated into leaves ectopically expressing an empty vector, wild-type or mutant PDLP5 as a pretreatment (Fig. 2B). Compared with empty vector control, *N. benthamiana* plants pretreated with wild-type PDLP5 exhibited delays in systemic viral movement, which was manifested by reduced levels of GFP fluorescence in systemic leaves. The plants pretreated with the ΔN or ΔC versions of PDLP5 also exhibited delays in viral movement similar to those pretreated with wild-type PDLP5 (Fig. 2B), indicating that they are functionally similar in this regard. By contrast, plants pretreated with Δ(N+C) lacking both signal sequences were similar to the empty vector control in regard to TMV-GFP movement. This result indicates that mislocalization is detrimental to PDLP5 function but that having one intact targeting signal is sufficient to confer function to PDLP5.

### PDLP family members all carry one or two targeting signals within JMe

Next, we investigated if other PDLP members are similar to PDLP5 in having two signals in the JMe sequence segment using a refined PdHMM (see Methods) followed by targeted mutagenesis. The decoding prediction suggested that all JMe sequences except the one derived from PDLP1 have two putative signal sequences (Table S1). Notably, they are all predicted to have one signal in a position consistent with that of N signal in PDLP5. Guided by the decoding results, we conducted targeted mutagenesis analyses against 6 PDLP members (Table 1) — note that we excluded PDLP2 from the test because it is highly similar (79% identity) to PDLP3.

**Table 1.**
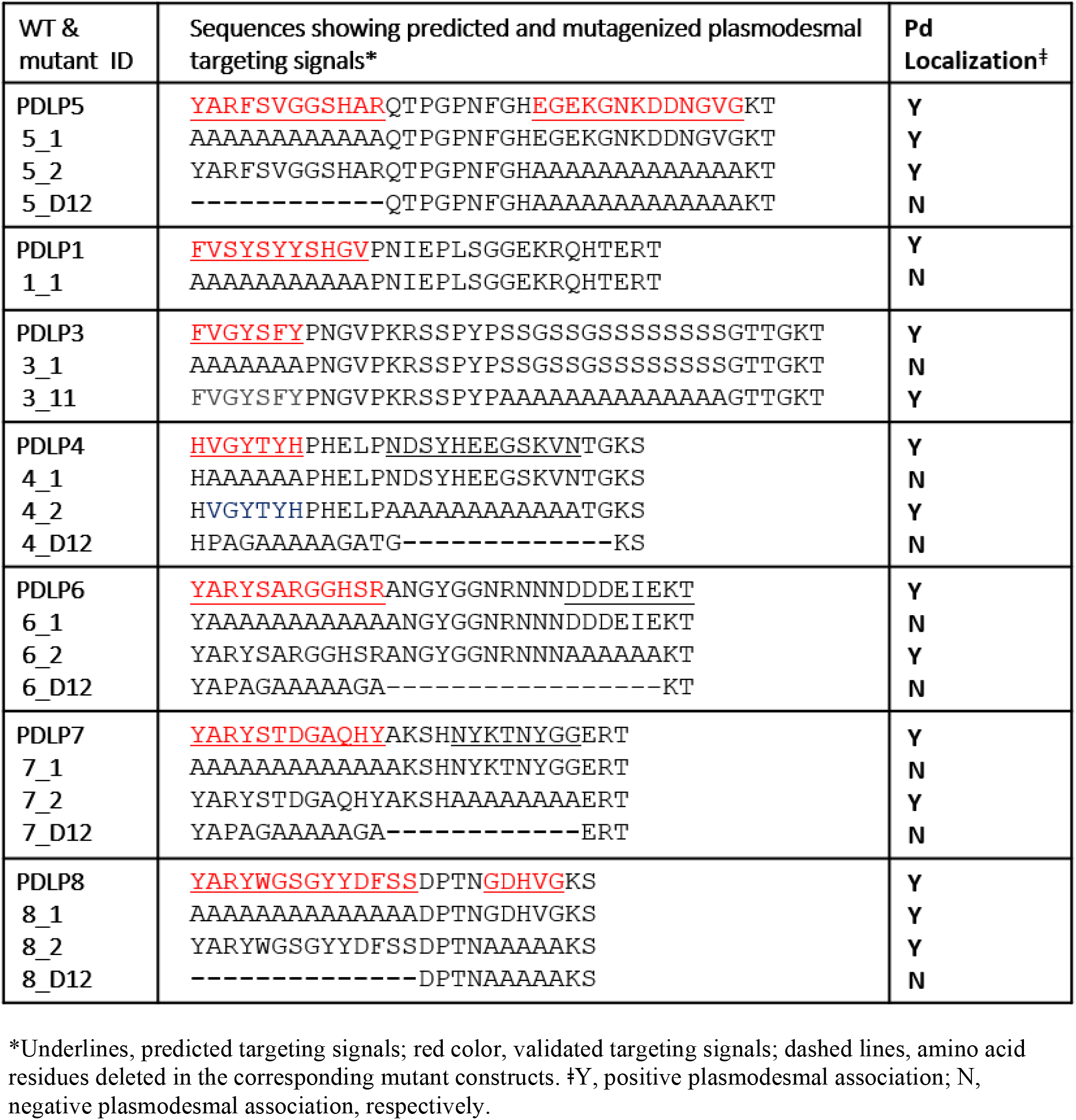
A summary of plasmodesmal targeting signals identified.

Subcellular localization studies of these mutated PDLP sequences revealed that the N-terminally positioned signals were all correctly identified vital for correct plasmodesmal localization (Table 1 & Fig. S7-S10). However, most of the C-terminally identified signals turned out to be dispensable for targeting. These include relatively long stretches of serine in PDLP3 (Fig. S8).

Intuitively, these repeated serine residues are unlikely to confer specificity as a signal necessary for recognition by a cargo carrier, and indeed they were dispensable for targeting. Lastly, the second signal predicted for PDLP8, which is relatively short, is indeed a true signal redundant to the N signal and thus similar to the case for PDLP5 (Fig. S10).

Collectively, these results demonstrate that PDLP members are similarly targeted to plasmodesmata via the signal(s) embedded in their JMe.

### Plasmodesmal targeting signals may be transferrable to heterologous proteins

To gain insight into how the targeting signals may function, we examined if the JMe sequence of PDLP5 could effectively target non-plasmodesmal proteins to plasmodesmata. Considering the structure of PDLP5, we chose the recipient proteins among type-I TM proteins and attempted to substitute a JMe equivalent region on those proteins with the PDLP5 JMe sequence. However, with this approach, we encountered problems that seemed related to protein folding and stability — potentially stemming from a disruption of the extended alpha-helical structure of TMDs by inserting JMe up close to the recipient TMD. Such an extended helix is predicted to be common in single-pass TM proteins (Lomize and Pogozheva, 2017). Therefore, we took a slightly modified approach by utilizing a plasmodesmata-association and anchoring cassette (PaaC) as a minimally functional unit required for targeting to and anchoring at the plasmodesmal membrane. The idea is to combine the three segments of JMe, TMD, and cytosolic tail into one molecular package for transfer.

Using the PaaC concept, we designed an experimental scheme, which we refer to as “PaaC swapping” (Fig. 3A). First, we assigned the TMD and extra- and intra-cellular JM sequences of a recipient protein of interest as a target region for swapping. Here, the JM sequences were manually assigned to be similar in length to those of PDLP5 PaaC (∼30-35 residues for the extracellular part and ∼5-10 residues for the intracellular part). To position JMe regions rationally, we considered where the ExD of the recipient protein might end or the cytosolic domain (e.g., a kinase domain) start if relevant information was available. It should be noted that this region exhibits no discernable sequence pattern across PDLPs or other proteins tested in our study. Next, as a test recipient protein, we chose BAK1, a well-known PM-localizing RLK. As expected, wild-type BAK1-EGFP localized to the PM quite evenly, exhibiting no trace of punctate fluorescent signals indicative of plasmodesmata. By contrast, an aberrant localization pattern was observed with BAK1 carrying PDLP5 PaaC. While still localizing to the PM, it showed a distinct plasmodesmal pattern, strongly overlapping with callose signals (Fig. 3B).

**Fig. 3.**
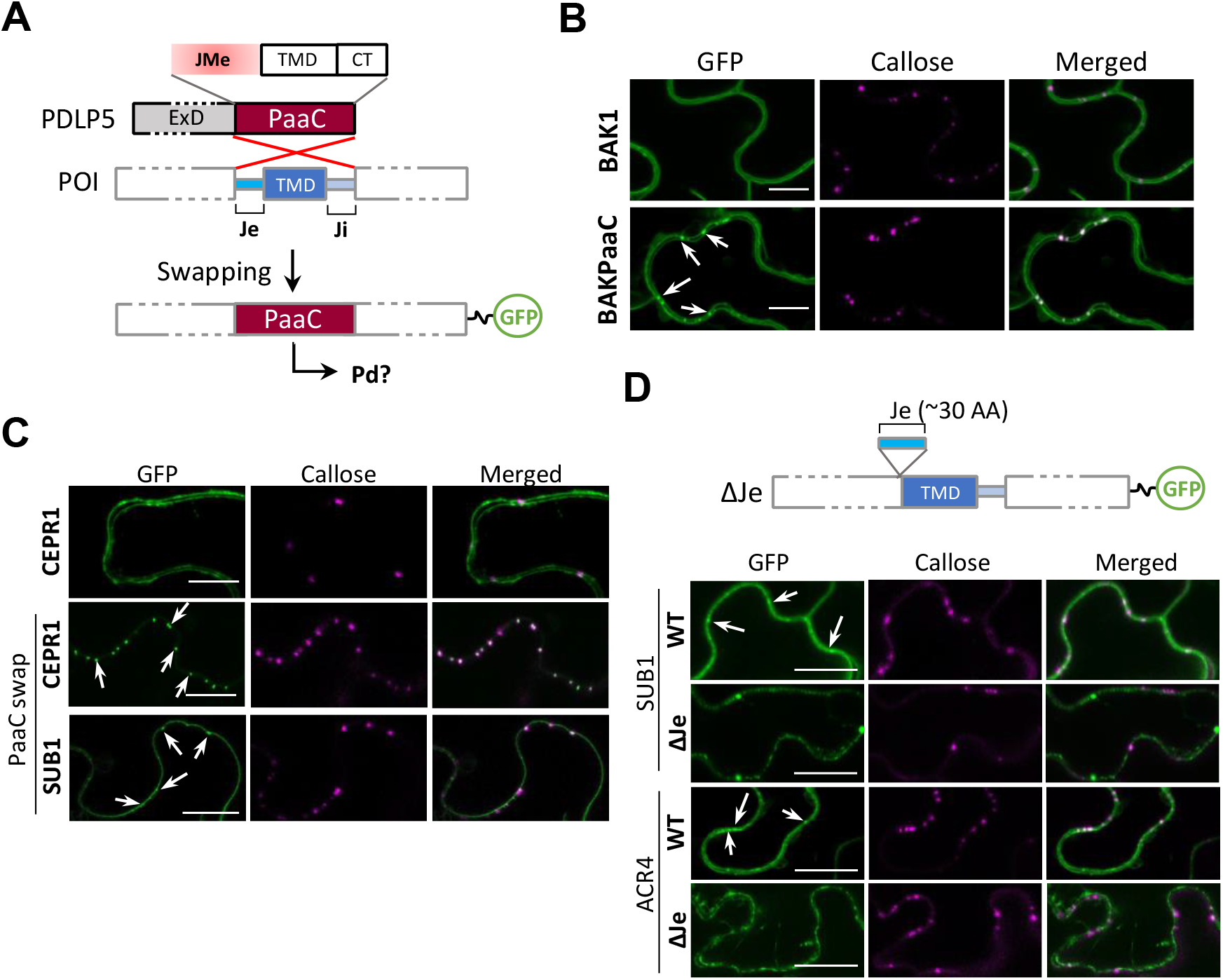
PDLP5 targeting signals are transferable. **A**. A diagram illustrating the experimental design of the PaaC-swapping mutagenesis to examine if the targeting signal in PDLP5 is transferrable. Je and Ji, regions comparable to JMe and CT, respectively, in PDLP5. **B**. Localization of wild-type BAK1 and the BAK1PaaC swap protein, each fused to EGFP. **C**. Localization of wild-type CEPR1 and its PaaC swap protein along with SUB1PaaC, each fused to EGFP. **D**. Localization of WT and mutated forms of SUB1 and ACR4, lacking the Je region corresponding to PDLP5JMe. Arrows, plasmodesmata. Scale bars, 10 µm.

This result suggests that PDLP5 PaaC is functional in a heterologous protein to redirect it to plasmodesmata.

Encouraged by this result, we then chose another non-plasmodesmal RLK, CEPR1, for PaaC swapping. Fluorescent signals of wild-type CEPR1-EGFP were predominantly associated with the PM and ER (Fig. 3C). In stark contrast, the presence of PDLP5 PaaC in CEPR1 redirected the protein to plasmodesmata. Next, we created a PaaC swap protein taking a plasmodesmal protein, SUB1, as a recipient. SUB1 has been shown to require neither its ExD nor cytosolic kinase domain but only its TMD for plasmodesmal association (Vaddepalli et al., 2014).

Excitingly, the resulting mutated SUB1 protein carrying PDLP5 PaaC in place of its corresponding native protein segment retained its plasmodesmal association (Fig. 3C). This result suggests that PDLP5 PaaC could also target SUB1 to plasmodesmata. These results add to the idea that PDLP5 PaaC may serve as a transferable targeting signal.

### Extracellular signal-based plasmodesmal targeting may apply broadly

That PDLP5 PaaC substitutes the native SUB1 TMD for plasmodesmal targeting led us to ask if extracellular signal-based plasmodesmal targeting serves as a common mechanism. We decided to test this possibility by examining if putative JMe regions have a role in subcellular localization in non-PDLP proteins such as SUB1 and ACR4, which are both RLKs associated with plasmodesmata but have unrelated ExDs. We assigned ∼30-aa residues as their putative JMe sequences for deletion mutagenesis, considering annotation information available from UniProt Knowledgebase (uniport.org). As expected, full-length SUB1 and ACR4 were both localized to the PM and plasmodesmata under our experimental conditions (Fig. 3 D). In contrast, lack of the putative JMe sequence altered their localization patterns, retaining the resulting mutated proteins in the ER and intracellular vesicles (Fig. 3 D).

Subsequently, we performed swap mutagenesis, in which PDLP5 in place of its native PaaC now had a PaaC-equivalent region derived from SUB1 or ACR4 (Fig. 4A). The resulting PDLP5 mutants were unaffected in plasmodesmal localization, indicating that SUB1 and ACR4 are likely to include functional PaaCs (Fig. 4B). We then performed the same test by choosing two additional plasmodesmal RLKs, FLS2 and BAM1. The resulting mutated PDLP5 proteins were again correctly targeted to plasmodesmata. These results support that these RLKs each likely carry a functional PaaC of their own. For further verification, we performed additional tests by including non-plasmodesmal RLKs, BAK1 and CEPR1, as donors of PaaC equivalent regions. In these cases the, mutated PDLP5 proteins carrying PaaC equivalents derived from these RLKs were impaired in plasmodesmal targeting and were instead misdirected to the PM (Fig. 4C).

**Fig. 4.**
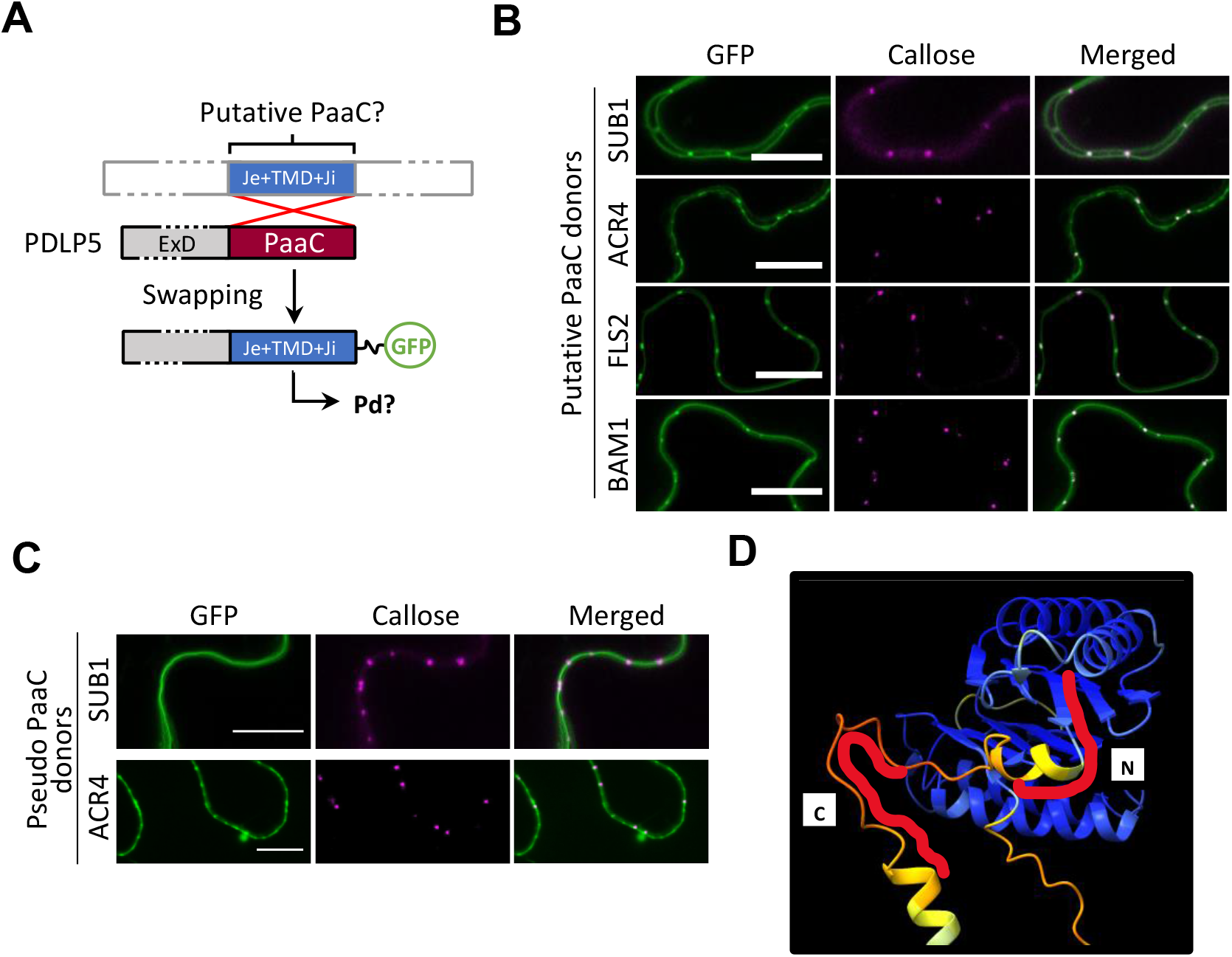
Extracellularly positioned targeting signals may be common. **A**. A diagram illustrating the experimental design to swap PDLP5 PaaC with a PaaC equivalent region derived from plasmodesmal or non-plasmodesmal proteins. **B**. PDLP5 carrying a putative PaaC region derived from four different plasmodesmal proteins. **C**. PDLP5 carrying a putative PaaC region derived from two non-plasmodesmal proteins. Scale bars in **B** & **C**, 10 µm. **D**. PDLP5 structure predicted using AlphaFold2 via ChimeraX with the JMe region in focus. Contoured red lines, positions of N and C signals.

These results underscore that PaaCs, likely via JMe regions, appear to serve as common targeting signals in a range of plasmodesmal proteins.

In this study, using a set of representative proteins, we have shown that plasmodesmal proteins require a stretch of amino acid residues serving as a targeting signal. It is a most striking finding that the signals are located in an extracellular region close to the membrane, which is unusual.

Notably, our experimental data point to the intriguing possibility that this extracellular signal-based plasmodesmal targeting may serve as a common mechanism among plasmodesmal targeted proteins.

## DISCUSSION

How plasmodesmal membrane proteins target plasmodesmata has been a long-standing question that has remained unresolved. In this study, we now show, using PDLP5 and representative single-pass plasmodesmal proteins, that plasmodesmal targeting requires a peptide signal encoded within the primary sequence of the proteins. This finding indicates that these proteins are delivered to plasmodesmata via specific transport machinery that recognizes those targeting signals. However, such machinery might also be novel because the carriers have to have specialized cognate receptors that are equipped with extracellular recognition domains. The majority of literature describes localization signals of membrane proteins being located in either cytosolic domains or within TMDs (Cosson et al., 2013; Barlowe and Helenius, 2016). Some animal membrane proteins that are targeted to the PM in a polarized manner are speculated to carry the targeting signals in their extracellular domains (Di Martino et al., 2019). To our knowledge, no plant proteins have been shown to utilize an extracellularly located targeting signal for their delivery to a specific membrane-bound compartment.

Interestingly, while the majority of PDLP members turn out to carry only one signal, PDLP5 consists of redundant targeting signals in tandem. These signals in PDLP5 are dissimilar to each other as well as to those in other PDLP members. Considering that mislocalization of PDLP5, caused by eliminating both targeting signals, impairs its function, specific localization of PDLP5 to plasmodesmata seems an essential prerequisite for its function. However, given that one targeting signal is sufficient for plasmodesmal localization and function of PDLP5, what advantage is there in having two signals? One possibility is that having two signals may serve as a mechanism to ensure that it is exclusively targeted to plasmodesmata, hence likely ensuring its potency. These two targeting signals may each constitute a strong signal with relatively high specificity and affinity to cognate receptors or add to the binding strength collectively. In this scenario, PDLP5’s association with non-plasmodesmal membrane would be unlikely. On the other hand, PDLPs with single targeting signals may potentially associate with the PM in addition to plasmodesmata, hence allowing them to exert dual functions. Indeed, the majority of plasmodesmal membrane proteins localize to the PM primarily and plasmodesmata seemingly secondarily. It would be interesting to find a correlation between having one signal and dual localization, such that a single or relatively weak signal allows for dual localization and hence, a dual function. Another possibility is that the two signals in PDLP5 might recruit independent receptors, hence reinforcing plasmodesmal localization. Hypothetically, these receptors might have differential affinities or differential expression patterns and levels depending on the physiological and developmental conditions of cells. In *Arabidopsis*, PDLP5 is expressed at very low levels but subjected to upregulation when plants become infected or undergo a specific developmental program in the root (Sager et al., 2020). Having two targeting signals may allow PDLP5 to target plasmodesmata in various cell types or cellular conditions.

Because our approach using an HMM-based machine learning algorithm is innately limited in terms of defining the core residues or the boundary residues, it is not clear how many and exactly which aa residues constitute the essential targeting signals. It is noteworthy that additional development of new computational methods is currently underway, which will allow in silico mutagenesis to pin down key residues using Fisher scores extracted from Pd-HMM and gradient descent. Nevertheless, PDLPs seem to be conserved in carrying one signal within the JMe region encompassing the last part of the second DUF26 module. This segment overlaps with the terminal β-sheet of the DUF26 protein fold (Vaattovaara et al., 2019b). In PDLP5, this side of the signal (N signal) pertains to 12-aa residues with the first half forming a β-sheet conserved in the DUF26 fold (Fig. 4D). The second half, together with the rest of the JMe sequence, including C signal, forms a low-confidence loop-like structure according to Alphafold prediciton. While the β-sheet forming part of the N signal is buried inside the DUF26 fold, making it unlikely to be available for binding, the remaining residues seem likely to be more accessible for binding, just as the signal C should be. Other members, PDLPs 1, 6, 7, and 8 have an N signal of similar length to that in PDLP5, which is thus likely accessible through its second half. In PDLPs 3 and 4, the signals end almost with the β-sheet. However, interestingly, the DUF26 folds in these members are more open than in PDLP5, suggesting a possibility that the targeting signals might still be accessible for binding. Another point to make here is that disrupting the β-sheet to facilitate targeting might impact protein folding. However, mutating this signal is unlikely to cause structural instability, given that mutating N signal does not impair protein localization or function of PDLP5 nor localization of PDLP8 (see Table 1).

Another striking feature of plasmodesmal targeting signals that we have identified in this current study is that they lack homologous sequences while being consistent with their location. Similar situations have been reported with certain animal receptors that undergo polarized protein sorting (Di Martino et al., 2019). Some receptors carry sorting signals that have completely different properties or sequences but occur in the same position, for example in the C-terminal tail.

However, the exact pathways and molecular components recognizing these unconventional sorting signals remain to be discovered. Similarly, with the plasmodesmal proteins that are the subject of this report, a lot more questions need to be answered in future investigations. Perhaps with the discovery of additional targeting signals in combination with new computational models, we may gain insight into what potential features the diverse signals may share. For example, more discernable sequence patterns may emerge as more examples become available to gain collective statistics. While being powerful enough to capture subtle patterns, HMMs are susceptible to the so-called black-box issue as most machine learning techniques are. In other words, the decoding outputs generated using Pd-HMM may not be readily interpretable as predicted signals do not reveal clear patterns as conserved motifs or sequences.

Having identified targeting signals, one of the most critical questions to address in future investigations will be how the signals are recognized for delivery to the final destination and how the proteins carrying the signals are loaded onto the plasmodesmal membrane. For targeting signal recognition, the putative plasmodesmal cargo receptors would have to have a cognate domain that is also extracellularly oriented, which would be novel. In terms of the delivery pathway, we can speculate one scenario in which plasmodesmal proteins are packaged into secretory vesicles and trafficked to the PM by default, followed by either lateral diffusion or an endocytic path to reach the plasmodesmal PM leaflet. Another scenario conceivable is that they may be targeted directly to plasmodesmata by departing from the early secretory pathway at some point post-ER or -Golgi to get onto a direct trafficking pathway to plasmodesmata. Now, with the discovery of plasmodesmal targeting signals, we are hopeful that other cases of this unconventional secretory pathway to plasmodesmata and associated novel trafficking machinery will be unveiled.

The operation of plasmodesmata is considered one of the mysteries of plant biology for we still do not know much about them at the molecular levels. Over the last few decades the field had seen tremendous progress towards understanding what molecules traffic through plasmodesmata and how plasmodesmal regulation is integrated with various cellular signaling pathways. In the current study, we have discovered the first few examples of plasmodesmal targeting signals, which have evaded scrutiny, in large part because of their attributes being unconventional, both in location and compositional diversity. Now, having these examples, should facilitate asking critical questions for future investigations such as the molecular composition of the targeting machinery and how pathogens might have exploited the innate system. We anticipate that an integrated approach, combining computational modeling and molecular and cellular experimentation, will reveal key aspects of protein targeting to plasmodesmata. In particular, this approach will be essential to identify hidden molecular patterns associated with the targeting signals and cognate receptors, which will eventually lead towards demystifying plasmodesmata.

## Materials and Methods

### Plant material and growth conditions

Wild-type *N. benthamiana* and *A. thaliana* (Col-0) plants were grown in a controlled environment under diurnal conditions of 18 hr light (120-180 μmol m^-2^ s^-1^) at 23°C and 6 hr dark at 21°C. The growth chambers were kept 24 hr at 60% relative humidity.

### Plasmid constructs

DNA constructs carrying mutations and various fusions were produced by overlapping PCR using a high-fidelity Taq Polymerase (Q5, New England Biolabs) followed by subcloning of gel-purified PCR fragments into a *Sfi*I-digested pMB binary vector (Sager et al., 2020) using T4 DNA ligase (New England Biolabs). To produce a C-terminal or an N-terminal fusion, pMB35S:*Sfi*I-EGFP and pMB35S:SP-citYFP-*Sfi*I were used, respectively. The latter plasmid is designed to carry at the N-terminus an SP (signal peptide) derived from PDLP5, to aid the cloning of type-I TM proteins. Also, both vector plasmids were designed to carry an alanine (APAGAAAAAGA) or a glycine (RPGGGGGP) linker between a protein of interest and either of fluorescent protein. The DNA sequence of each plasmid construct was confirmed for its fidelity using Sanger sequencing. The amino acid sequence information of PaaC or PaaC equivalent regions are provided in Table S2 and the nucleotide sequence information of all primers used to create plasmid constructs used in this study is provided in Table S3.

### Agrobacteria-mediated gene expression

*Agrobacterium tumefaciens* cells of strain GV3101 (+pSoup) were transformed with each pMB plasmid harboring a specific DNA construct by electroporation. Transformants were selected by culturing the cells overnight at 28°C on LB medium supplemented with gentamicin 50 μg/ml, rifampicin 50 μg/ml, tetracyclin 10 μg/ml, and spectinomycin 200 μg/ml. Positive colonies were grown in liquid LB media containing the same antibiotics overnight at 28°C by shaking at 225 rpm. Harvested cells were resuspended and diluted to the desired optical density (OD) (0.4 at 600 nm) in infiltration buffer comprising 10 mM MES, pH 5.7, 10 mM MgSO_4_, and 100 µM acetosyringone. Resuspended agrobacterial cells were used to infiltrate mature leaves of 3-to 4-week-old *N. benthamiana* plants. Following agroinfiltration, plants were kept in a growth chamber for an additional 2-4 days until they were imaged by microscopy. Subcellular localization studies were performed using at least 2-3 plants per construct and repeated at least three independent times.

### Live-cell imaging and processing

Subcellular localization was performed using a Zeiss LSM 880 or Leica Stellaris 8 tauSTED confocal microscope. Leaf segments were prepared for examination by mounting them in water and using single well Lab-Tek®II Chambers with coverslips. Leaf cells were imaged using a C-Apochromat X40/1.20-W Korr UV-VIS-IR or 86x/1.2W MotCorr objective with the 488 nm Argon laser and 505-550 nm band-pass emission filter for EGFP signals and 514 nm excitation line and 517-579 nm emission for YFP signals. Detection of aniline-blue stained callose was performed using the 405 nm Diode laser and a 420-480 nm band-pass emission filter. Images were acquired as small Z-stacks of optical sections and rendered as 3-D projections using ImageJ.

### Viral movement assays

Two mature leaves (4th and 5th) were first infiltrated with agrobacterial cells transformed with pMB35S vectors carrying no insert (empty vector) or wild-type or mutated forms of PDLP5.

Three days later, the same leaves were secondarily infiltrated with agrobacterial cells transformed with a binary vector carrying a recombinant TMV genome encoding free GFP. The plants were monitored for systemic viral movement for the next several days. For the primary infiltration, agrobacterial cells carrying no insert or wild-type or mutated PDLP5 were diluted in the infiltration buffer to OD 0.8. The cells carrying the viral suppressor protein p19 were diluted to OD 0.6. Thus prepared, agrobacterial cells were mixed in equal volumes before infiltration of target leaves. Agrobacterial cells carrying TMV-GFP were diluted to OD 0.02. Plants were imaged under UV illumination through a deep yellow filter mounted on a Nikon D3100 digital camera. Viral movement assays were performed using at least 5 plants per treatment and repeated at least three times.

### Computational modeling, decoding predictions, and structural visualization

To build an HMM architecture we formulated two assumptions; one, that the JMe consists of two signals, and two, that the two signals may be separated by a non-signal residue(s) (Li et al., 2020a). Thus, the resulting Pd-HMM1.0 designed to capture plasmodesmal targeting signals is a three-state HMM. In addition, to build our model with limited unlabeled training examples, in addition to the standard Baum-Welch algorithm, we developed two novel algorithms (Li et al., 2020b; Li et al., 2021) to enable active learning and the use of both positive training examples (PDLPs) and negative training examples (non PDLPs). For decoding of the JMe sequence of PDLP5, PdHMM1.0 was designed as a 3-state hidden Markov model using the Baum-Welch algorithm, having an architecture as detailed elsewhere (Li et al., 2020a).

To train the model, JMe sequences of PDLPs1-8 and the 10 best hits of PDLP5’s orthologues were used. Following validation of the two targeting signals in PDLP5, we refined the model to version 2.0. This version was trained with a novel HMM training algorithm that modifies the Baum-Welch algorithm so that partial label information can be used for better performance, as demonstrated elsewhere (Li et al., 2021). In general, there are two standard training algorithms, one using fully labeled examples (Maximum Likelihood) and the other using fully unlabeled examples (Baum-Welch, aka Expectation-Maximization). To upgrade PdHMM, we developed a novel training algorithm that enables active learning and is capable of training an HMM with partial labels, which in our case are PDLP5’s confirmed targeting signals (Li et al., 2021).

To visualize the predicted structure of the JMe region, a full-length PDLP5 aa sequence was submitted to the AlphaFold2 (Tunyasuvunakool et al., 2021) program through the software UCSF ChimeraX (Pettersen et al., 2021) according to the user guide.

## ACKNOWLEDGMENTS

We thank S.J. Streatfield for thorough editing.

## Author contributions

Conceptualization: JYL

Methodology: J-YL, GRL, WX, LL, JL

Investigation: GRL, JL, J-YL, LL, WX

Visualization: GRL, J-YL

Funding acquisition: J-YL, LL

Project administration: J-YL, LL

Writing – original draft: J-YL

Writing – review & editing: J-YL, GRL, JL, LL, XW

## Data and materials availability

Materials are available from the corresponding author upon reasonable request.

## SUPPLEMENTARY MATERIALS

Materials and Methods Table S1 - S3

Fig S1 – S10

References (23 – 28)

